# Immunogenicity studies of various experimental vaccines in chickens

**DOI:** 10.1101/2020.08.31.276154

**Authors:** Angel Justiz Vaillant, Patrick E. Akpaka

## Abstract

In this paper, the main objective was to raise chickens’ antibodies against three crucial public health microorganisms: the human immunodeficiency virus-1, *Salmonella* spp, and *Staphylococcus aureus*. Immunogens were prepared from the said microorganisms. Chickens were vaccinated either orally or intramuscularly. After a booster immunization, mostly eggs were collected and assess for the presence of specific antibodies. The most important results were the production of a large amount of anti-HIV antibodies in chicken’s eggs, and also the synthesis of anti-protein A antibodies with the ability to inhibit the growth of *S. aureus in vitro* and to serve as anti-anti-idiotypic antibodies with the capacity of neutralizing the original antigen. Enzyme-linked immune absorbent assays detected the presence of these antibodies as anti-Salmonella antibodies that were critical in reducing the bacterial load in the stomach and caeca compared with a control group. The vaccines were effective and safe, but more laboratory work, and economics have to be carried out to start a human trial.

## Introduction

Purifying antibodies called immunoglobulin Y (IgY) isolated from the egg yolk of chickens is of particular interest as a source of specific antibodies for oral administration to prevent infections and to use them as a reagent in immunodiagnostic procedures. The use of birds in antibody production results in a reduction in the use of laboratory animals. Immunized chickens produce larger quantities of antibodies (2000 mg IgY/month) than do rodents (200 mg IgG/month) in the laboratory [1].

According to Jerne's network theory, it is possible to produce an antibody against another antibody's antigen-binding site [2]. In this paper, we tested two hypotheses. The first hypothesis was that the immunization with viral and bacterial products (immunogens) can shed a potent immune response that can be assessed in the eggs of chickens. The immunogens used were from human immunodeficiency virus (HIV), *Salmonella* spp, and *Staphylococcus aureus*. The second hypothesis was that an orally administered antibody stimulates the production of a complementary antibody so-called anti-idiotypic, which can be used therapeutically. It has been an avenue for vaccine development for decades; however, it has not given the expected result. We also aimed to raise anti-*Salmonella* antibodies in chickens, which successfully reduced the bacterial load in the stomach and caeca of immunized birds as compared with an unvaccinated control group of birds, where full colonization of bacteria was confirmed.

## Materials and methods

Ethical approval was granted from the Campus Ethics Committee of the University of the West Indies, Mona campus. The laboratory work described was carried out under the EU Directive 2010/63/EU for experiments.

A higher confidence level entails a greater sample size. Power is the probability of statistically significant evidence of a disparity among the groups, given the population's variation. Higher power requires a larger sample size. The SpA [3] and HIV vaccine [4] study previously published had a small animal number. This new experiment that will be reported here aimed to report previous findings of the development of specific antibodies using oral hyperimmune egg as feeding. It proved useful in the case of the vaccine against S. aureus [3] and a pilot study of an oral vaccine against HIV, using cats fed chicken eggs [4]. However, it reduced the production of anti-HIV antibodies in cats' blood, when viral synthetic peptides were used as immunogens. Still, the animal size was too small, and we need to repeat the experiments using a different approach [4]. The use of 3 anti-HIV intramuscular vaccines and testing the chicken eggs, rather than the blood of domestic cats, was used in this second approach [2].

In the case of the Salmonella vaccine in our first published study, we evaluated two vaccines, a DNA vaccine and a live-attenuated vaccine [5]. This last one gave the expected results and was economical in its preparation. The amount of birds used in this experiment is representative of the broiler chicken population in Jamaica. Due to how vital is the poultry industry on the island, and the challenges it has faced [6], we wanted to have a statistical representation justified by the number of birds used.

The chicken house is made of wood and recreates the natural habitat of chickens. It has a black shade around the house that provides 70% shading. The area to the sides and front of the chicken house was weeded and cleaned for esthetics and minimizing bacterial growth. The shaded house helped reduce the admission of sun rays, which, coupled with the zinc roof, would increase the chicken house's temperature, which is undesirable for the birds. Chickens were kept in separate cages. Egg collection was performed daily, including weekends.

Chicken house material consisted of Hypo Layer feed, 1.5m gauged mesh, 0.25 gauge expanding metal, and push broom shower. Other elements include water buckets, bleach (sodium hypochlorite), hydrogen peroxide 5%, egg trays, industrial glove, sterile latex glove, sterile applicator sticks, 27G1/2 size syringe, feed tray, and water trays.

The chickens were tested and observed for four months or another specified period. No dewormer, vitamins, or medication of any sort was administrated to the birds under the project. The animal house was washed out with cleaning products twice daily, and fecal content removed in keeping with a sound biosecurity practice. The chickens were monitored for well-being and health daily and supervised by a veterinary technician. Fresh water and corn were given to the birds at free demand. Chicks were fed with anti-SpA hyper-immune egg and water freely as a part of the S. aureus vaccine project [3].

The chicken mortality rate was zero percent in the both groups, in the intramuscular immunized (IMI) as well as in the orally immunized (OI). Five birds become sick after being vaccinated with the live-attenuated *Salmonella* vaccine via orally, but they recuperated themself in a few days. In this group weakness, and decreased appetite were the most common symptoms that accounted for 16.6% (5/30), but yellow diarrhea was observed in one bird which represented 3.33% (1/30). No antibiotics were administered to them. The Salmonella vaccine being researched is made of various strains of the bacteria (sensitive strains to much antibiotics), that proved to be safe in a pilot study where seven chicken were inoculated with no apparent side effects [5]. We noticed a decrease in egg production 3 to 7 days after immunization. We did not realize that in the pilot study. Nevertheless, probabilistically, in the more extensive clinical study, some side effects may become more apparent since there is a more significant number of birds. Indeed, It justifies the use of a greater sample size of animals for performing more extensive research and evaluate the actual impact in the use of different biologicals on birds.

### Immunogenicity studies of experimental HIV vaccine in chickens

All grade reagents used in this study were commercially available. They were purchased from Sigma-Aldrich. The studies were repeated three times, and it was obtained similar results. The HIV immunogens used in these experiments were the keyhole limpet hemocyanin (KLH) conjugated to HIV peptides (fragments 308-331 and 421-438 of the HIV gp120) and (fragment 579-601 of the HIV gp41) [2].

The amino acid sequences of HIV peptides and their references are cited below.

HIV-gp41 (579-601): Arg-lle-Leu-Ala-Val-Glu-Arg-Tyr-Leu-Lys-Asp-Gln-Gln-Leu-Leu-Gly-lle-Trp-Gly-Cys-Ser-Gly-Lys [7].

HIV-gp120 (308-331): Asn-Asn-Thr-Arg-Lys-Ser-lle-Arg-lle-Gln-Arg-Gly-Pro-Gly-Arg-Ala-Phe-Val-Thr-lle-Gly-Lys-lle-Gly [8].

HIV gp120 (421-438): Lys-Gln-Phe-lle-Asn-Met-Trp-Gln-Glu-Val-Gly-Lys-Ala-Met-Tyr-Ala-Pro-Pro [9].

### Dimerization of HIV peptides (addition of a C-terminal cysteine) and preparation of HIV immunogens

The C-terminal cysteine was added to the amino acidic sequences of HIV peptides (fragment 579-601 of the HIV-gp41 and fragments 308-331 and 421-438 of the HIV-gp120). These peptide fragments were dimerized by cysteine oxidation with dimethyl-sulfoxide [10]. Each HIV peptide was dissolved in 5% acetic acid to a final concentration of 5.1 mg/ml. The pH of the medium was adjusted to 6 with 1 M (NH_4_)2CO3, and dimethyl-sulfoxide was added to 20% of the final volume, and after four hours at room temperature (RT), the solute was extracted. After that, the peptide was dissolved in 3 ml 5% trifluoroacetic acid and precipitated with 35 ml cold ether. The precipitate was dialyzed against 1.2 liters of deionized water, pH 7 at 4°C overnight [2].

Then, 1 mg of keyhole limpet hemocyanin (KLH) was diluted in 2.1 ml 0.1 M borate buffer (1.24 g boric acid, 1.90 g sodium tetraborate, pH 10, in 500 mL deionized water). In a 20 ml glass tube, with a gentle stirring, 1.1 µmol of each HIV synthetic peptide (with C-terminal cysteine added) and 0.22 milliliters 0.3% glutaraldehyde solution (ACS reagent grade, pH 5.5, Sigma-Aldrich) at RT were slowly mixed and left to stand for 1.50 hrs. A yellow coloration was observed (this indicated that the conjugation process was successful). To blocking the excess of glutaraldehyde, 0.26 ml of 1 M glycine (Sigma-Aldrich) was added, and the mix was left for 32 min at RT. Each HIV-hemocynin conjugate was then dialyzed against 1.3 liters 0.1 M of borate buffer, pH 8.4 through the night at 4°C, then the same buffer was used to dialyze the preparations for 8 hrs at 4°C. The dialysates were stored at 4°C until further use [2].

### Chicken immunization

Six healthy brown Leghorn layer hens (two per HIV immunogen), aged seven months, were immunized intramuscularly (IM) at multiple sites on the breast with a specific KLH-conjugated HIV peptide vaccine. Chickens were vaccinated on day 0, with 0.5 mg/ml of the immunogens in ml of complete Freund’s adjuvant (Sigma-Aldrich), and on days 14, 28, and 45 after the first immunization, hens received booster doses of 0.25 mg/ml of the immunogen in 0.5 ml incomplete Freund’s adjuvant. The eggs were collected daily pre- and post-immunization. Additionally, 15 healthy brown Leghorn were immunized, six birds per HIV vaccine with a similar immunization schedule as above. Eggs were collected on day zero and 45 days after to evaluate the reproducibility of the ELISAs for determination of anti-HIV antibodies. The coefficient of variation (CV) intra-assay and inter-assay were calculated for each one of the vaccines.

The water-soluble fraction (WSF) contains an elevated IgY concentration, and it was separated from the lipid content by the partial application of the Polson methodology [2], which uses chloroform only, as follows:

1. Wash the egg with warm water and crack the eggshell carefully.
2. Separate the egg yolk from the egg white manually and place the egg content on a tissue paper to help to remove as much egg white as possible.
3. Wash four times the egg yolk with 100 ml of PBS, pH 7.4 (Sigma-Aldrich).
4. Break the egg yolk membrane and pour it into a 50 ml tube. Then, dilute egg yolk 1:5 in PBS, pH 7.4, and add an equal volume of chloroform (ACS reagent grade, Sigma-Aldrich).
5. Shake the mixture, and vortex and roll the tube containing the mixture on a rolling mixer for 15 minutes.
6. Centrifuge the mixture for 15 min (2000×g at 18°C).
7. Decant the supernatant, which contains the water-soluble fraction (WSF) that is the diluted egg yolk plasma, rich in IgY (25 mg/ml), which is approximately four times the IgY concentration present in the serum (6 mg/ml).

### Indirect enzyme-linked immunosorbent assay (ELISA for the investigation of the presence of anti-HIV antibodies in the WSF of chickens)

#### Preparation of ELISA reagents

##### Coating buffer

3.7 g Sodium Bicarbonate (NaHCO_3_) and 0.64 g Sodium Carbonate (Na_2_CO_3_) in 1L of distilled water.

##### Phosphate buffered-saline Tween-20 (10% PBS-Tween 20, pH 7.2)

Dissolve the following: 0.2g of KCl, 8g of NaCl, 1.45g of Na_2_HPO_4_, 0.25g of KH_2_PO_4_, and 2ml of tween-20 in 800 ml of distilled water. After that, adjust pH to 7.2, add additional distilled water to adjust the volume to 1L, and then may sterilize by autoclaving.

##### Blocking solution

It is used to prevent or stop the non-specific binding of antibodies and antigens to the microtiter well. Preparation is as followed: add 0.1 g KCl, 0.1 g K_3_PO_4_, 1.16 g Na_2_HPO_4_, and 4.0 g NaCl to 500 ml distilled water, pH 7.4. Then, to complete the preparation of this solution, 15g of non-fat dry milk should be added.

##### Sample/Conjugate Diluent

Add 15 g of non-fat dry milk and 2.5 ml of 10% Tween 20 to 500 ml of PBS.

#### ELISA for anti-HIV peptide antibodies

The 96-well polystyrene microplates (U-shaped bottom, Sigma-Aldrich) were coated with 100 ng of 579-601 of the HIV gp41, fragments 308-331 or 421-438 from HIV gp120 in coating buffer for 4.1 h at 37°C. Then, each microplate was washed four times with 10% PBS-Tween 20 and the blocking solution (3% non-fat milk in PBS) added in the amount of 51 μl into each well. The microplates were incubated at 1.30 h at RT. After that, microplates were washed as previously. Fifty μl of WSF diluted 1:50 with the sample diluent was added to the wells. Then, each microplate was incubated for 1h at RT and washed four times as previously. After that, 50 μl of horseradish peroxidase labeled anti-IgY conjugate (Sigma-Aldrich) diluted 1:30,000 was poured into each well. Microtiter plates were incubated again for 1h at RT and washed four times. A volume of 50 μl of tetramethylbenzidine (TMB, Sigma-Aldrich) was added; and after a further incubation of 16 min in the dark, the reaction was stopped with a solution of 3M HCl and each microplate was read in a microplate reader at 450 nm. The cut-off value was assessed from the mean optical density (OD) of the negative control times 2. The cut-off points of ELISAs for the detection of anti-HIV peptide (579-601), anti-HIV peptide (308-331) and anti-HIV peptide (421-438) were 0.42, 0.40 and 0.44 respectively [2].

Positive and negative controls were homemade. Four positive controls were used in each assay. They were prepared from egg yolk samples with the highest titers of specific anti-HIV peptide antibodies and their OD values between 1.20 and 1.50 at 450 nm. Four negative controls were used in each assay. They were prepared from the egg yolks of non-immunized animals, and they showed OD values of 0.170-0.20 at 450 nm. Three replicates of each WSF sample per bird, collected on day 60 were assayed for the presence of anti-HIV peptide antibodies using the present ELISA

Table 1 shows the results of immunogenicity studies of an HIV vaccine in brown Leghorn layer hens.

**Table 1.** Results of immunogenicity studies of experimental HIV vaccine in brown Leghorn layer hens.

| ELISA results of the 3 experimental vaccines | Mean optical density (XOD)<br>Pre-immunization (day 0) | Mean optical density (XOD)<br>Post-immunization (day 45) | P-value |
| --- | --- | --- | --- |
| HIV-gp41 (579-601) | 0.170 (0.021) | 0.885 (0.044) | P<0.01 |
| HIV-gp120 (308-331) | 0.156 (0.015) | 0.910 (0.024) | P<0.01 |
| HIV-gp41 (421-438) | 0.188 (0.01) | 0.865 (0.038) | P<0.01 |

### Immunogenicity studies of a Salmonellosis experimental vaccine in brown Leghorn layer hens

#### *Salmonella* strains: life-attenuated vaccine preparation and immunization protocol

The immunogen used to protect against salmonellosis in brown Leghorn layer hens contains five live-attenuated serovars of *Salmonella* strains isolated from poultry products and environmental samples: *Salmonella* Montevideo, *Salmonella* Yeerongpilly, *Salmonella* Augustenborg, *Salmonella* Kentucky and *Salmonella* Typhimurium. To many, 30 birds were given intramuscularly (at multiple sites in the breast) 10^8^ CFU/ml of a mix of *Salmonella* strains (immunogen) contained in 1 ml of incomplete Freund’s adjuvant (IFA) at day 0 and on days 14^th^ and 28^th^ after the first immunization, hens received booster doses of 10^5^ CFU/ml of the mix. The said immunogen without IFA was given to another group of 30 birds per-orally in a quantity of 10^3^ CFU/ml contained in 1 ml of 25% non-fat milk at day 0, 14, and 28. A control group of 30 birds has injected 1 ml of 0.9% saline solution containing 0.4 ml of IFA also at multiple sites of the breasts at day 0, 14, and 28. The WSF of egg yolks in the three groups (intramuscularly immunized (IMI), orally immunized (OI), and control group (CG) was prepared from yolks of eggs collected before and after the immunotherapy (40-42 days post-immunization). Triplicates of WSF (180 samples) were assayed for the presence of anti-*Salmonella* antibodies using the ELISA described below [5].

### Isolation of Salmonella from Specimens

The *Salmonella* Typhimurium isolation was carried out as followed: the exterior of the hen cloaca was first cleaned with a sterilized and moistened cotton balls before application of the cotton tips of each swab applicator. The swabs, *caeca*, and stomach tissues were immediately placed in a sterile screw-cap test tube containing 9 ml of pre-enrichment broth (buffered peptone water 1%). At least 2.5 g of each type of specimen was dissolved in 250 ml pre-enrichment broth. The inoculated pre-enrichment broth was incubated at 37 °C for 24 hrs following this incubation it was thoroughly mixed using a vortex mixer. A 1.1 ml aliquot of buffered peptone water 1% was added to 9 ml of enrichment broth (selenite broth, selenite cystine broth, and tetrathionate broth) and further incubated at 37 °C for 24 hrs. After vortexing 0.16 ml and a 3 mm loopful of inoculum was used to inoculate the differential plating media such as *Salmonella Shigella* agar selective media, MacConkey agar, bismuth sulphite agar and brilliant green agar that were incubated at 37 °C for 24-48 hrs. Following the incubation, as typically, the cultures were examined, and non-lactose fermenting colonies were selected and used to inoculate Kleiger iron agar and urea agar slants. After a further 24 hours incubation period at 37°C colonies that gave the typical *Salmonella Shigella* reaction, were inoculated to the routine line of sugars and again incubated. Confirmation was followed by slide agglutination with somatic “O” and flagella “H” antigens of *Salmonella*. Serological typing of *Salmonella* Typhimurium was performed [5,9].

### Identification by slide agglutination

Presumption *Salmonella* Typhimurium isolates were stored on tryptose agar a room temperature until confirmation as previously described (Kauffman-White Schema, Difco Laboratory, Detroit, and Michigan U.S.A). For each isolate, 2 loops of the growth on tryptose agar was emulsified in 1 drop of normal saline solution (0.9%) on a microscope slide. The preparation was examined for autoagglutination. If the organism was not self-agglutinating, one drop of either “H” or “O” antiserum was added to each spot. The slide was agitated by gently rocking back and forth for 2 to 3 minutes after mixing. The slide was examined for agglutination [4].

### Antibiotic Susceptibility Test

All Salmonella isolates tested were investigated for their antibiotic resistance with the disc diffusion test using the following discs (Difco): gentamicin (10 μg), kanamycin (30 μg), ampicillin (10 μg), amikacin (30 μg), trimethoprim/sulfamethoxazole (1.25/23.75 μg), chloramphenicol (30 μg), cefazolin (30 μg), cephalothin (30 μg), cefepime (30 μg), cefotaxime (30 μg), streptomycin (10 μg), ceftazidime (30 μg), cefoxitin (30 μg), nalidixic acid (30 μg), ciprofloxacin (5 μg), norfloxacin (10 μg), tetracycline (30 μg) and imipenem (10 μg) [4].

### Purification of *Salmonella*

A 1:9 *Salmonella* suspension was made in buffered peptone water and incubated overnight at 37°C. One ml of pre-enrichment broth was transferred to a 1.5 ml micro-centrifuge tube and centrifuged for ten minutes at 14,000 × g (Eppendorf Model 5424), the supernatant was carefully discarded. The pellets were re-suspended in 300 μl sterile PCR grade water by vortexing. The tube was again centrifuged at 14,000 × g for five minutes. The supernatant was discarded with care. The pellets were again resuspended in 300 μl PCR grade water by vortexing. The microcentrifuge tube was incubated at 100°C for 15 minutes and immediately chilled on ice. The tube was centrifuged at 14,000 × g at 4°C. The supernatant was transferred to a new tube and incubated at 10 min at 100°C then chilled immediately on ice. The supernatant was stored at −20 °C [3,11].

### Immunoglobulin-Y isolation

The chicken immunoglobulin Y (IgY) fraction was isolated by the chloroform-polyethylene glycol (PEG) method [12].

### Enzyme-linked immunosorbent assay (ELISA) for studying the presence of anti-Salmonella antibody in layer hens

U-shaped bottom's ninety-six well polystyrene microplates purchased at Sigma-Aldrich, St. Louis USA were incubated with (2 μg/well) of the LPS (Sigma –Aldrich) from *Salmonella* Typhimurium in coating buffer (overnight at 4 °C.) The microtiter plates were washed four times, with 10 % PBS-Tween-20 and blocked with 3% non-fat milk in PBS (25 μl/well) and incubated 1 hr at RT. The microplates were washed four times. Then a 50 μl aliquot of the previously isolated egg yolk (Ig)Y solutions in a concentration of 1.25 mg/ml was added in triplicate. After incubating for one hour at RT, the microplates were washed four times, and 50 μl of the anti-IgY-HRP conjugate (Sigma-Aldrich) diluted to 1:30000 with conjugate diluent (prepared as previously described in the preparation of ELISA reagent section) was added into each well. Microplates were incubated for 1 hr at RT. Then, they were washed four times and 50 μl tetramethylbenzidine (TMB, Sigma-Aldrich) was added into each well. Microplates were further incubated for 15 minutes in the dark, and 50 μl 3M HCl was added to stop the reaction. After that, reaction color development was measured with a microplate reader (Synergy™ Neo Hybrid Multi-Mode Microplate Reader). The cut-off point was an OD of 0.51, and it was calculated from the XOD of the negative control times 3 [5]. Table 2 and 3 show results of immunogenicity study and *Salmonella* contamination percentage in caeca and stomachs after challenge with the wild typed S. Typhimurium, respectively. The ELISA tested triplicates of a total of 90 IgY preparations.

**Table 2.**
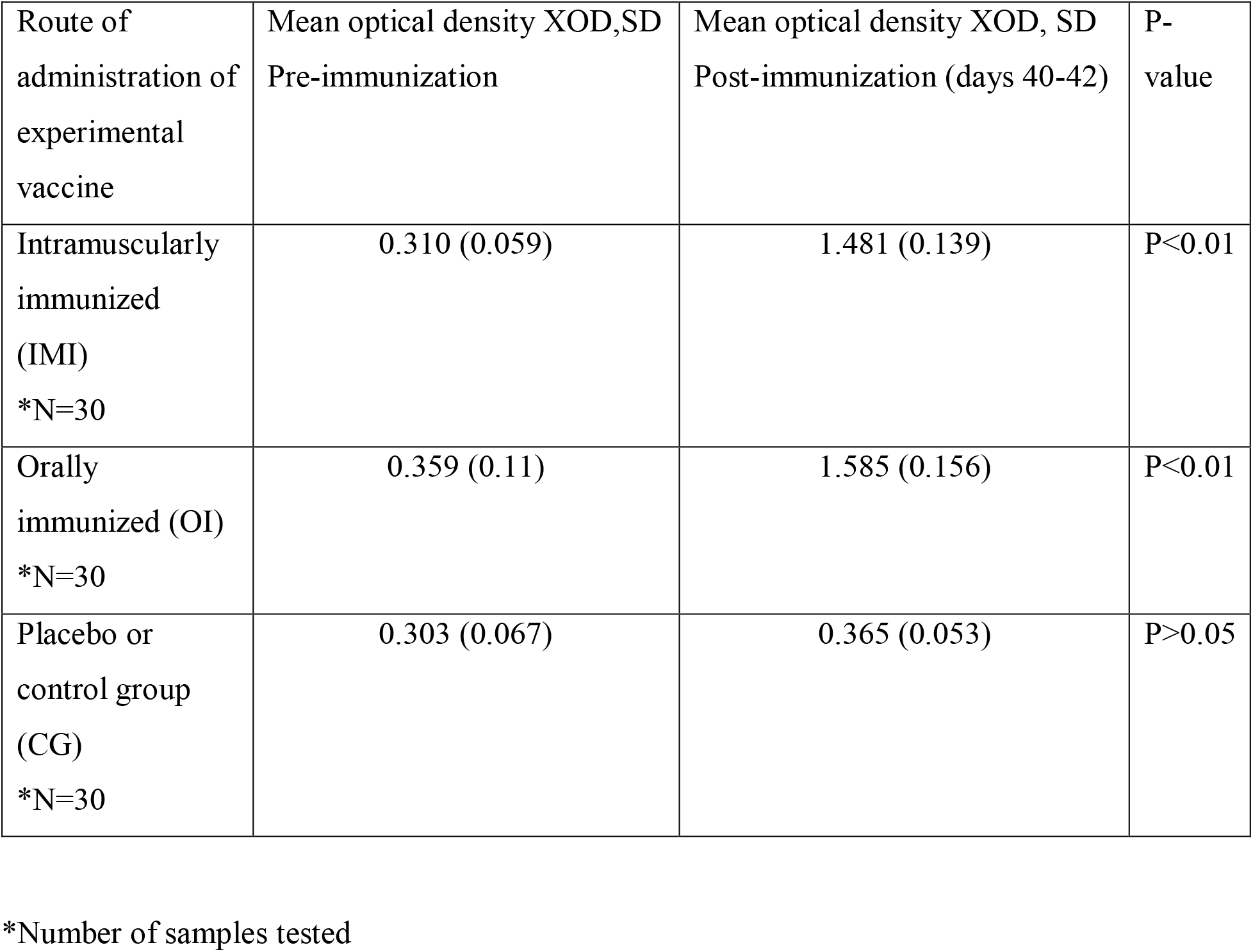
Results of immunogenicity studies of experimental Salmonellosis vaccine in brown Leghorn layer hens.

| Route of administration of experimental vaccine | Mean optical density XOD,SD<br>Pre-immunization | Mean optical density XOD, SD<br>Post-immunization (days 40-42) | P-value |
| --- | --- | --- | --- |
| Intramuscularly immunized (IMI)<br>*N=30 | 0.310 (0.059) | 1.481 (0.139) | P<0.01 |
| Orally immunized (OI)<br>*N=30 | 0.359 (0.11) | 1.585 (0.156) | P<0.01 |
| Placebo or control group (CG)<br>*N=30 | 0.303 (0.067) | 0.365 (0.053) | P>0.05 |

\*Number of samples tested

**Table 3.** *Salmonella* contamination percentage in caeca and stomach after challenge with the wild typed S. Typhimurium.

| Vaccinated birds and controls | Mean Concentration of *Salmonella* in caeca | Mean Concentration of *Salmonella* in stomach |
| --- | --- | --- |
| Live-attenuated vaccine (30 birds) | $10^2$ CFU/g | $10^2$ CFU/g |
| Control group (30 birds) | $2 \times 10^7$ CFU/g | $10^5$ CFU/g |
| P-value | $P<0.01$ | $P<0.01$ |

The reproducibility of the ELISA for anti-Salmonella antibodies was assessed by running 12 positive and 11 negative samples in three different occasion, on the same day, and on alternative days. The coefficients of variation percentage were calculated as a measure of reproducibility.

### Experimental infection with a wild-type Salmonella Typhimurium

An experimental infection challenge was carried out forty-five days post-immunization to assessing if specific antibody production can stop or modify a *Salmonella* Typhimurium infection *in vivo*. Previously immunized birds (IMI and OI) and the control group (CG) a total of 90 animals were given via crop gavage an infected cocktail (1.25 ml containing 103 CFU/g equivalent to 50% lethal dose) of a wild multi-resistant-type of *Salmonella* Typhimurium, resistant to gentamicin, kanamycin, ampicillin, chloramphenicol, streptomycin, nalidixic acid, ciprofloxacin, norfloxacin, and tetracycline. Chickens were euthanized using 40% carbon dioxide with 30% oxygen and 30% nitrogen, and their anatomical organs were removed and placed in sterile bags to be analyzed for the presence of *Salmonella* spp [5], which in chickens may cause gastroenteritis in young birds. However, adult brown Leghorn layer hens can serve as lifetime hosts for this pathogen without showing signs of infection.

Table 3 shows the *Salmonella* contamination percentage in *caeca* and stomach after challenge with the wild type *Salmonella* Typhimurium. Immunized hens cleared the infection as their immune system reacted by raising the levels of specific antibodies to combat the microorganism. Despite *Salmonella* infection, five of the unprotected birds (treated with placebo) were visibly sick, but the infection rate of *Salmonellosis* in this population was 100%.

The rate of infection by organs showed that caeca were much predisposed to infection. It was previously reported [5,11]. As a natural mechanism of defense, the gut immune system produces specific IgA against bacteria that are not a part of the healthy flora. Somehow in chronic carriers of salmonellosis as adult birds, a mechanism of immunological tolerance operates and makes birds to ignore the presence of *Salmonella* spp that can co-exist in symbiosis with regular residents of the gut microbiota. It could explain the apparent resistance of adult birds to salmonellosis and their lack of clinical manifestations [5,6].

### Statistical analysis

It was done by using SPSS software (version 18). Differences between cases and controls were tested. A P-value<0.05 was statistically significant.

### Immunogenicity studies of an experimental vaccine against S. aureus in chickens

#### Protocol of the immunization of hens with Staphylococcal protein A (SpA)

Two healthy layer chickens (brown Leghorn), aged seven months, were injected IM on the breasts with 0.6 mg of Staphylococcal protein A in 0.5 ml complete Freund’s adjuvant (CFA) on day 0 and 0.25 mg of the same antigen in incomplete Freund’s adjuvant (IFA) on days 14, 28 and 42. The eggs were collected post-immunization. The water-soluble fraction (WSF) from the egg yolk, which is rich in antibodies IgY, was separated by the partial application of the method of Polson. ELISAs measured the presence of a range of anti-protein A antibodies in the WSF [4,12].

#### Feeding of chicks with hyperimmune eggs (containing Ab1 and Ab-2)

Hyperimmune eggs with a titer of 1: 3000 diluted ¼ volumes of cow’s milk were fed on demand to eighteen (18) chicks aged zero, divided into three groups. The first group was fed for one week, the second group for two weeks, and the third one was fed for four weeks. Another six chicks were fed non-hyper-immune egg solutions diluted in ¼ volumes of cow’s milk for four weeks. A 0.8 ml blood samples were taken at the end of the feeding program (at 8 days, at 15 days and at 4 weeks) [3].

#### Sandwich ELISA for detection of anti-SpA antibodies (Ab-1 and Ab-3)

U-shaped bottom microplates were coated with 500 ng of SpA (Sigma-Aldrich) in a coating buffer for four hours at 37°C. After four washing procedures with 50 μl/well of 10% PBS-Tween-20 and blockage with 3% non-fat milk in PBS, 25 μl/well, 1h at RT. The microplates were rewashed four times. Microplates were added 50 μl of WSF diluted 1:50 and prepared from the egg yolks of pre-immunized and post-immunized layer hens. A total of 32 samples were assayed for the detection of Ab-1. Alternatively, 50 μl/well of chick serum was added for the detection of Ab-3 (18 samples in total). After incubation for 1h at RT, the microplates were rewashed four times, and 50 μl of horseradish peroxidase-Staphylococcal protein-A conjugate (Sigma-Aldrich) diluted 1:3000 was added. After incubation for 1h at RT, microplates were washed and a TMB solution (50 μl) was poured to each well. After 16 minute's incubation in the dark, the reaction was stopped with 3M HCl and read in a microplate reader at 450 nm [3]. The cut-off point was 0.615 for Ab-1 and 0.586 for Ab-2. They were calculated as 3x XOD of the negative control.

### Purification by affinity chromatography of anti-anti-SpA antibodies (Ab-3) raised in the sera of chicks daily fed anti-SpA hyper-immune eggs

A commercial protein-A affinity chromatography called PURE-1A (Sigma-Aldrich.) was used to purify anti-anti-SpA antibodies (Ab-3) from sera of chicks daily fed (orally immunized) anti-SpA hyper-immune yolks. The instructions of the manufacturer were followed in performing this procedure. Briefly-Anti-SpA containing serum is first loaded onto the Protein A Cartridge where the IgG is immobilized. The Protein A Cartridge is then washed to remove excess unbound proteins. The Desalting Cartridge is readied for use by reactivating with HEPES buffer. The Protein A Cartridge and Desalting Cartridge are then connected via the Luer lock fittings and the Elution Buffer is introduced. The eluate contains the purified IgG at a physiological pH. Both cartridges may be regenerated and stored for future use [3].

A similar purification procedure was used to purify anti-SpA (Ab-1) present in undiluted fractions (1 ml) of WSF of egg yolks from hens immunized with SpA. Aliquots of IgY preparations (Ab-1 and Ab-3) were collected in 1 ml micro-tubes and kept at −20°C until further use [3]. The protein concentration of purified IgY (Ab-1 and Ab-3) was assessed by a commercially available ELISA kit (MyBiosource, Inc), which protocol was performed following the manufacturer’s instructions.

### Purification of anti-anti-SpA antibody (Ab-2) using SpA-bearing *Staphylococcus aureus* cells

A novel method was designed for the purification of Ab-2 from undiluted WSF of hyperimmune eggs. A total of six preparations were made. The procedure is as follows:

1. Mix in an Eppendorf microtube 0.9 ml of undiluted WSF from immunized birds with 25 μl of SpA-bearing *Staphylococcus aureus* cells (Sigma-Aldrich).
2. Incubate the microtube at 37°C for 30 min.
3. After the incubation period, centrifuge the microtube in an Eppendorf 5424 centrifuge for 5 min.
4. Observe the microtube that presents a pellet of Ab-1 bound-cells at the bottom, and the supernatant, where Ab-2 is present.
5. Decant the supernatant to another microtube.
6. Store it at −20°C until further use.

### Competitive ELISA to study the inhibition of the SpA-binding to Ab-3 by Ab-2

This immunoassay tested the ability of anti-anti-SpA (Ab-2) to inhibit or interfere with the binding of anti-anti-anti-SpA (Ab-3) to the original antigen-staphylococcal protein A (SpA). It is a confirmation test of the functional capacity of Ab-2. In this ELISA, both Ab-2 and the antigen SpA compete for binding to Ab-3. When Ab-2 is present, it blocks the binding of HRP-labeled SpA to Ab-3, resulting in inhibition of reaction color development. The procedure is as follows:

1. Prepare materials and ELISA buffer solutions and reagents as described above.
2. Pipette 53 μl of purified Ab-3 (100 μg/ml) mixed with 2.8 ml of coating buffer into each well.
3. Incubate the microplate at 37°C for 4 hrs.
4. Aspirate the contents of the wells.
5. Fill each well with an appropriately diluted washing solution and aspirate. Wash three times more.
6. Pipette 53 μl of serial dilutions of pooled Ab-2 (100 μg/ml) in triplicates.
7. Incubate the microplate at RT for 1 hr.
8. Rewash the microplate filling each well with 100 μl of washing buffer.
9. Pipette 53 μl of commercially available peroxidase labeled SpA conjugate (Sigma-Aldrich) diluted 1:5000 to each well.
10. Reincubate the microplate at RT for 1 hr.
11. Repeat step 5.
12. Pipette 53 μl of TMB (Sigma-Aldrich) to each well.
13. Incubate in the dark at RT for 14 min.
14. Measure absorbance at 450 nm in a microplate reader
15. Analyze the results.

The Ab-2 inhibition percentage of the Ab-3 binding to SpA was calculated using the formula:

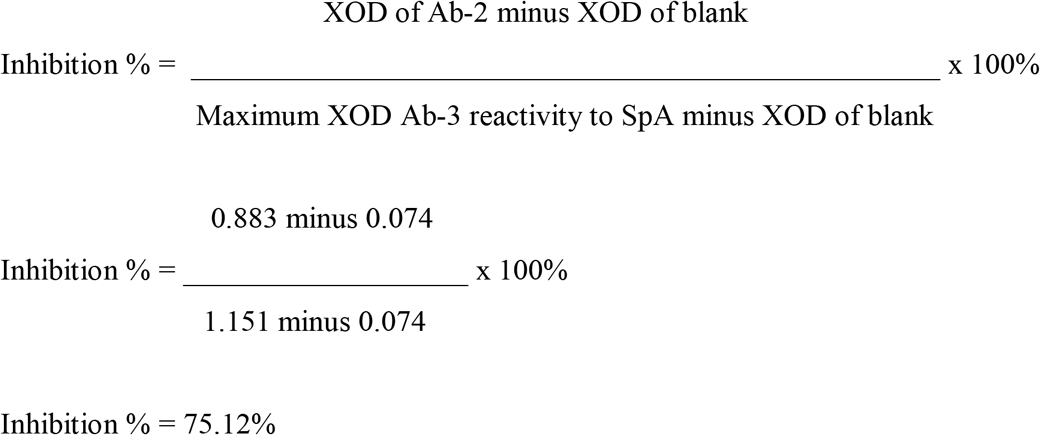

Note: this result suggests that the pooled Ab-2 interfered 75.12% with the inhibition of the specific Ab-3 binding to SpA. Besides, it represents a confirmation *in vitro* of the interaction that exists between the two idiotypic antibodies. Six samples were pooled to make the Ab-2 used in this assay.

### Study of the *in vitro* effect of purified serum anti-anti-anti-SpA antibodies (Ab-3) on cultures of *Staphylococcus aureus*

The investigation of the neutralizing ability of purified anti-SpA antibodies was carried out as follows: One ml of brain heart infusion (BHI) broths was placed in 10 sterile test tubes. To each was added an equal volume of 10 μl of purified anti-SpA antibodies in a concentration of 1.25-μg/μl. The amounts of samples tested were 18 samples from chicks fed with HIE, and six samples from chicks fed non-hyperimmune eggs. Replicates of a sample of 9% normal saline were used as a blank and added to tubes 11 and 12. All samples were tested in duplicate. An inoculum of an ATCC *Staphylococcus aureus* strain (ATCC #33592) was prepared to 0.5 commercially prepared McFarland scale standard (1=300 × 10^6^/ml bacteria concentration). Then, ten μl of this inoculum was added serially from tubes 1 to 10 (prepared for each sample) and the dilutions were reported in this study as a bacterial concentration of 0.1, 0.01, and 0.001. The absorbance value of each tube at 600 nm was measured. A sample from each tube was plated out on Blood agar and incubated overnight at 35°C. Inhibition of the bacterial growth was observed in individual and pooled sera of chicks fed with anti-SpA hyperimmune eggs up to 28 days, 15 days, and eight days period. Bacterial growth was observed when blanks or sera of chicks fed non-HIE were used because these samples lacked the protective Ab-3. The XOD value of the samples was plotted against the different bacterial concentrations at various serial dilutions [3], as shown in Figure 1–3.

**Figure 1.**
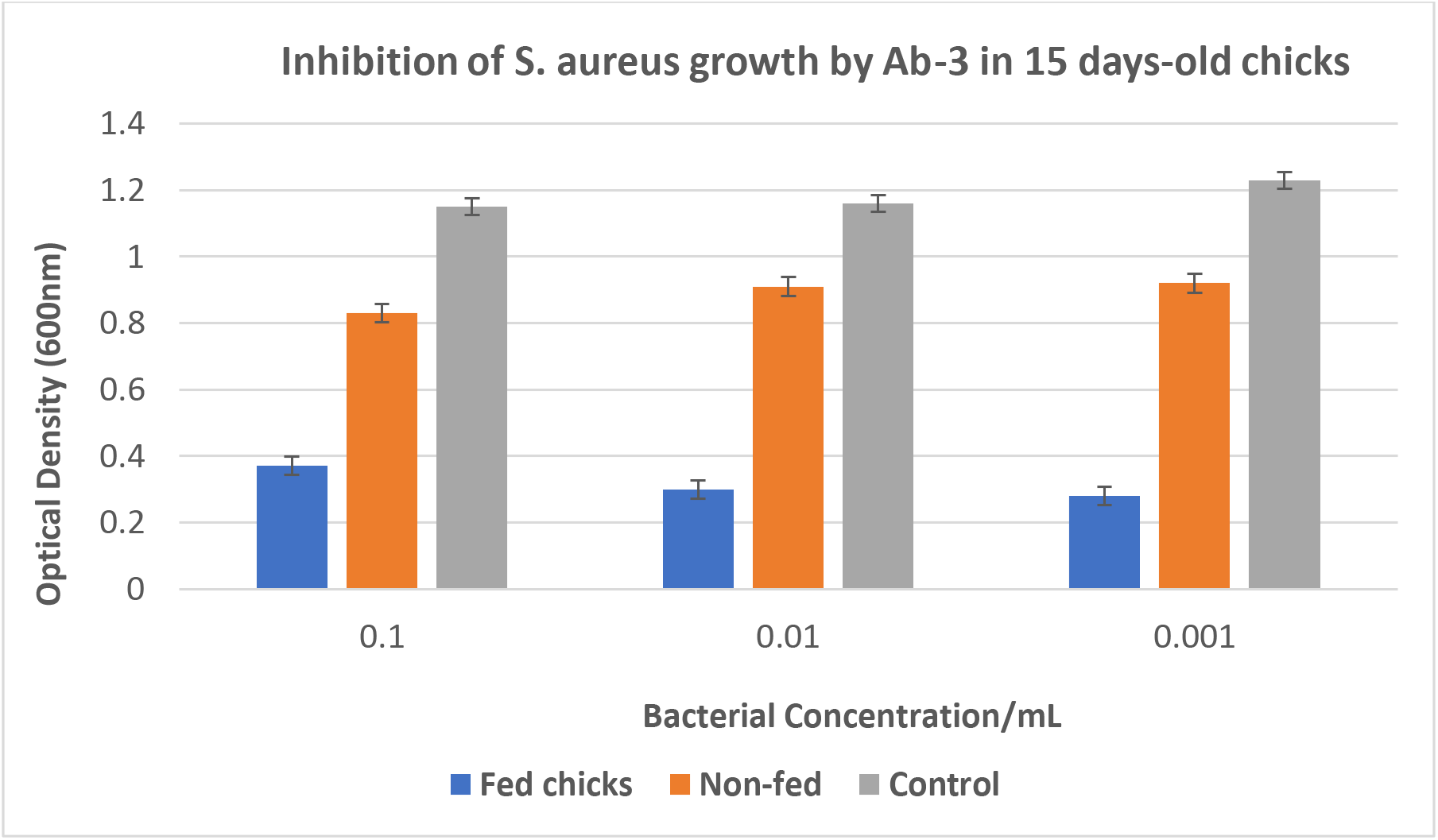
Inhibition of S. aureus growth by Ab-3 purified by SpA-affinity chromatography and observed in pooled sera of chicks (six) fed with anti-SpA hyperimmune eggs up to 15 days old. Inhibition was observed in dilutions of S. aureus treated, suggesting that the Ab-3 is a mirror image of SpA, binding to it on the cell wall of S. aureus inhibits the pathogen growth, it does not happen in non-fed chicks and controls. *Control: bacterial suspension without Ab-3. Non-fed animal: fed with non-hyperimmune eggs (eluates from purification, which tested negative for the presence of Ab-3 by the sandwich ELISA.

**Figure 2.**
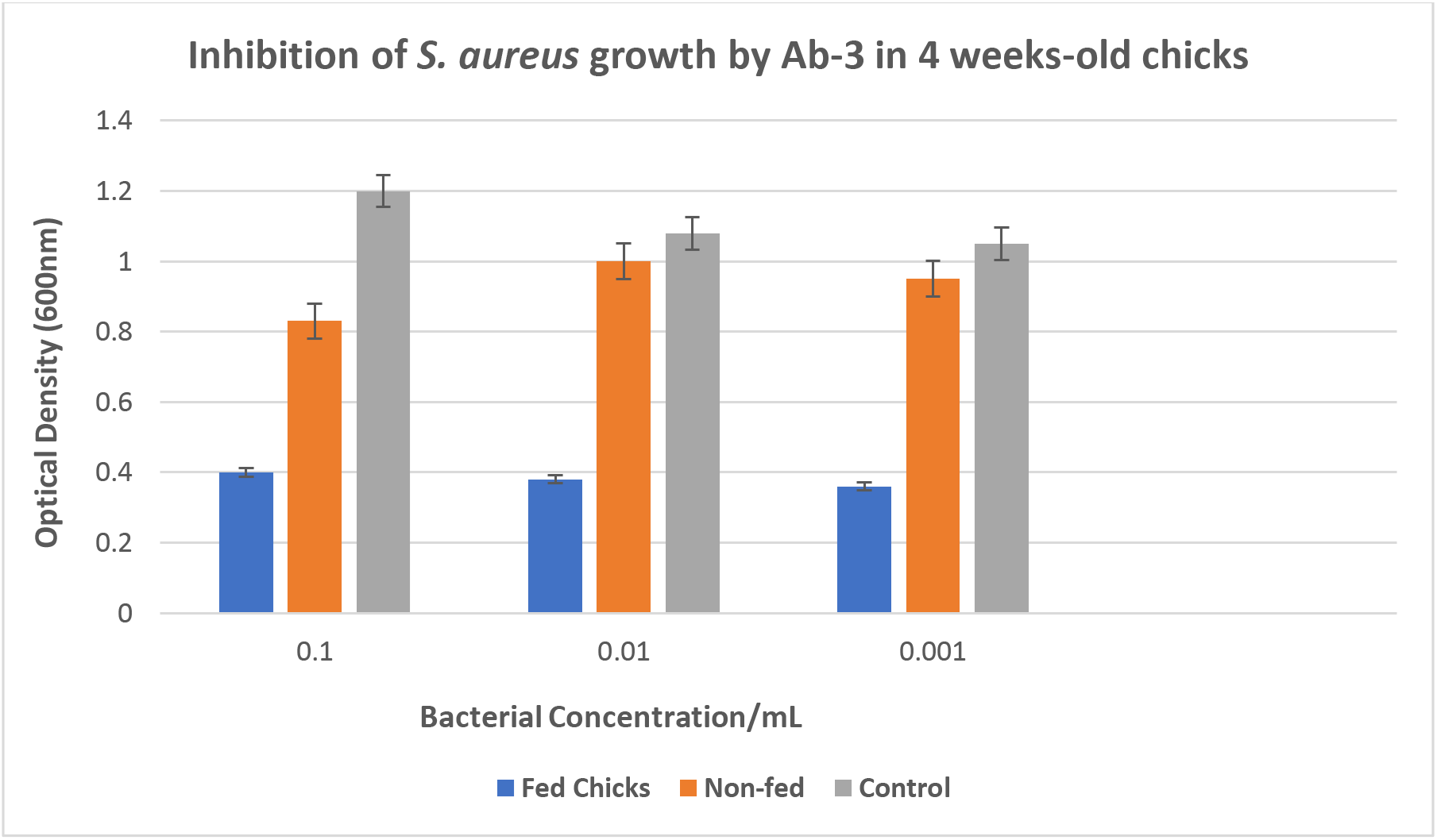
Inhibition of S. aureus growth by Ab-3 purified by SpA-affinity chromatography and observed in pooled sera of chicks (six) fed with anti-SpA hyperimmune eggs up to 4 weeks. Inhibition was observed in dilutions of S. aureus treated, suggesting that the Ab-3 is a mirror image of SpA, binding to it on the cell wall of S. aureus inhibits the pathogen growth, it does not happen in non-fed chicks and controls.

**Figure 3.**
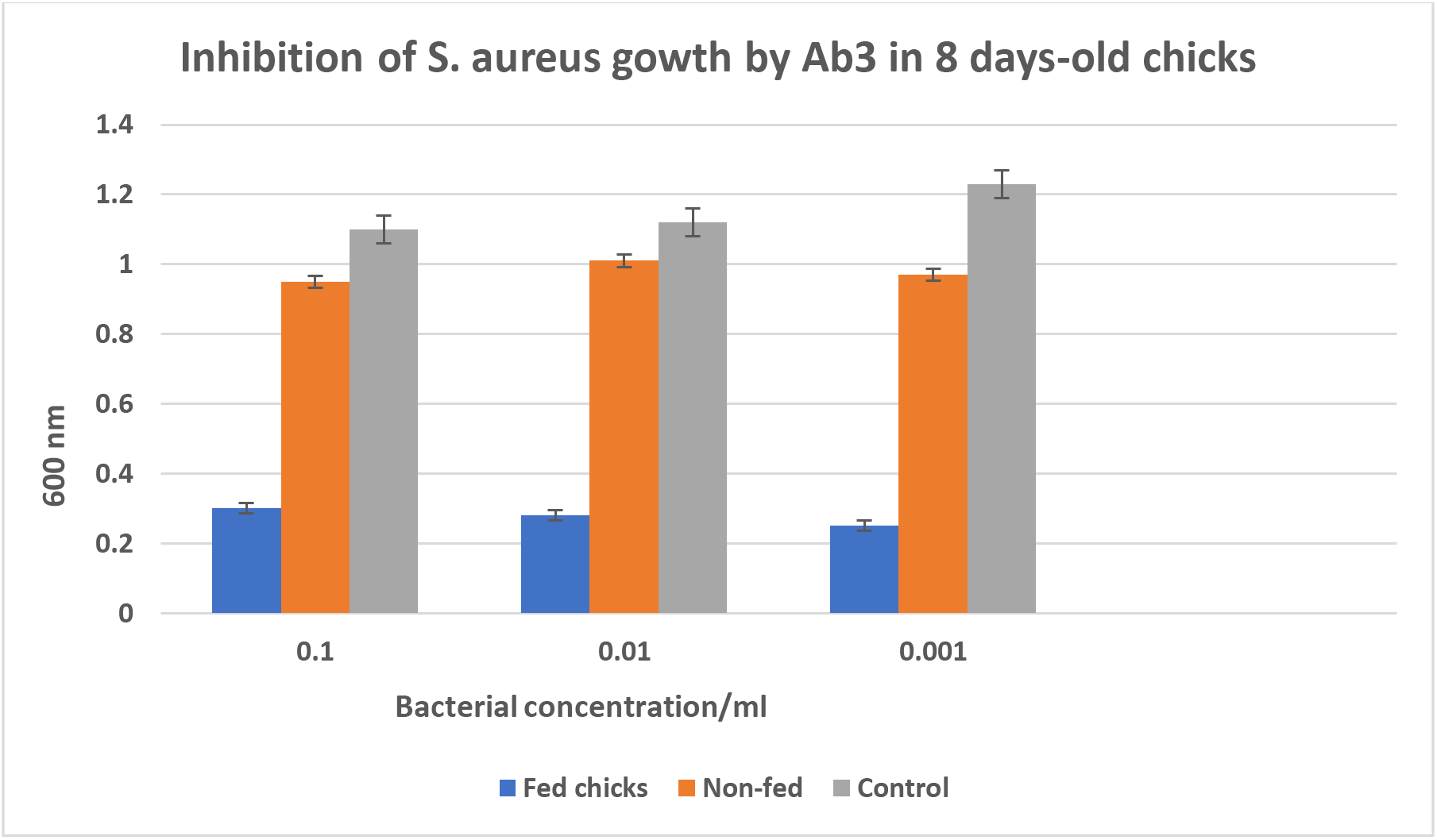
Inhibition of *S. aureu*s growth by Ab-3 purified by SpA-affinity chromatography and observed in pooled sera of chicks (six) fed with anti-SpA hyper-immune eggs up to 8 days. Inhibition was observed in dilutions of *S. aureus* treated, suggesting that the Ab-3 is a mirror image of SpA, binding to it on the cell wall of *S. aureus* inhibits the pathogen growth, it does not happen in non-fed chicks and controls.

### Statistical analysis

Fisher’s exact test (Epi Info 3.5.3 software, CDC, Atlanta, GA, USA) and the chi-squared test were used to compare the concentrations of the Staphylococcus aureus that are potentially inhibited by the groups of chicks that were the fed with anti-Staphylococcal protein-A antibody (anti-SpA Ab3) over the non-fed chicks (the OD value vs. the serial bacterial concentrations). The result was descriptive and was reported as a comparison of frequency distributions. A P-value <0.05 was considered significant.

## Results and discussion

Table 1 shows water-soluble fractions (WSF) of egg yolks collected zero- and 60-days post-immunization were assayed for the presence of anti-HIV peptide antibodies using ELISA (six samples in total were assayed and tested three time). These immunogenicity results showed that HIV peptide vaccines effectively produced strong anti-HIV immune responses in immunized brown Leghorn layer hens. There was a statistical level of significance (P<0.01) that explains the notable degree of difference (related to the anti-HIV antibody levels in the egg yolks) between pre-immunized and post-immunized birds in the three experimental vaccines [2]. These immunogenicity results showed that HIV peptide vaccines effectively produced a strong anti-HIV immune response in immunized brown Leghorn layer hens.

The results shows six healthy brown Leghorn layer hens (two birds per HIV vaccine), aged seven months that were immunized intramuscularly (IM) at multiple sites on breast with 3 specific KLH-conjugated HIV peptide vaccine (five birds per vaccine) on day 0, 15, 30 and 45. Eggs from pre-immunized and post-immunized birds were collected. The water-soluble fraction of each egg was isolated by few steps of the Polson method (1990) and tested by different ELISAs. The coefficient of variation intra- and inter-assay were within the normal limits. It suggests that the assays were reproducible (reported in supportive information). We could not compare our HIV tests with a commercially prepared assay because at the time of this study was not available and it is still not available.

Table 2 shows an enzyme-linked immunosorbent assay (ELISA) for anti-*Salmonella* antibodies, which assesses the immune response of Brown Leghorn layer hens immunized intramuscularly (IMI) and orally (OI) against *Salmonella* spp. A placebo or control group was included and studied as birds may present positive titers of anti-*Salmonella* antibodies constitutionally, as this pathogen can be found ubiquitously in chickens. The ELISA tested triplicates of a total of 90 IgY preparations (from egg yolks) [12]. This study demonstrated that immunized birds either IMI or OI had raised high titers of specific antibodies against antigens of various *Salmonella* serovars that cross-react with the lipopolysaccharide (LPS) of *Salmonella* Typhimurium [5]. There is a statistical significance that suggests a critical difference concerning the specific level of antibodies before and after immunization with the attenuated vaccine. The cut-off point was calculated as XOD of negative control plus 2 SD.

Eggs were collected pre- and post-immunization and the egg yolk proteins assessed for the presence of anti-Salmonella antibodies. The coefficient of variation (intra- and inter-assay) were within the normal limit, less than 5% for intra-assays and less than 10% for inter-assays.

Table 3 shows the challenge with the wild type *Salmonella* Typhimurium, where both immunized groups (IMI and OI) showed, 8-day post-challenge, a mean concentration of *Salmonella* in *caeca* and stomach statistically significantly different (P<0.01) from that of the control group, which showed a higher level of *Salmonella* colonization in these organs, since they were not immunized. The explanation remains in the presence of protective anti-*Salmonella* antibodies in vaccinated birds that fights off *Salmonella* infection. On the other hand, the control group that was injected 1ml of 0.9% saline solution in incomplete Freund’s adjuvant did not exhibit immunity against salmonellosis as it indeed lacked specific antibodies.

The results shown here presents that polyclonal antibodies anti-HIV, anti-*Salmonella*, and anti-*S. aureus* had in their paratopes the information to create anti-ID antibodies (Ab-2) able to release anti-anti-ID (Ab-3). This was first described by Niels Jerne more than 40 years ago and still nowadays it is hopefully used to produce new vaccines (much experimental) against pathogens [13–17]. The mechanism of the generation of Jerne’s network is as followed: An antigen (immunogen) stimulates the synthesis of Abs (Ab1). The active centers of a molecule Ab1 are recognized by the second Abs—anti-IDs (Ab2). This last one (Ab2) serves as an antigen or immunogen for another third class of Abs (Ab3)—anti-anti-IDs, and so on. The Jerne’s network appeared to be unfinished or endless, because Ab3 and Ab1 are closely and molecularly related and show identical or nearly recognizing properties among antibodies [13]. This paper demonstrates that was possible to produce an idiotypic SpA antibody (Ab-1), anti-ID SpA (Ab2), anti-anti-ID SpA (Ab3) to protect against *S. aureus in vitro.*

These Ab-3 were able to neutralize the original antigen: SpA. For instance, Ab-3 developed against S. aureus was able to inhibit the growth of S. aureus *in vitro* and to inhibit bacterial plaque formation. This approach could be an avenue for treating bacterial infections caused by S. aureus, including respiratory and skin infections in other species. It may try in humans.

On the other hand, immunization of young chickens with *Salmonella* antigens could protect birds from Salmonellosis. We hypothesize that this protection will pass on to immature birds since extremely high titers of anti-Salmonella antibody are present in the egg yolk, to protect the embryos and later the chicks [18].

To demonstrate the hypothesis that eggs can be a good model to study ID-anti-ID interactions we immunize two healthy layer chickens (brown Leghorn), aged seven months. They were immunized IM on the breasts with 0.6 mg of Staphylococcal protein A in 0.5 ml complete Freund’s adjuvant (CFA) on day 0 and 0.25 mg of the same antigen in incomplete Freund’s adjuvant (IFA) on days 3000 diluted ¼ volumes of cow’s milk were fed on demand to eighteen (18) chicks aged zero, divided into three groups. Once the chicks grew, blood was taken from six chicks after seven days, and another two groups of six chicks at 2 weeks and 4 weeks, respectively. Ab-3 were purified using a SpA-affinity chromatography system (Sigma-Aldrich Co). These antibodies were able to neutralize the growth of *S.aureus* in vitro by inhibiting bacterial colony formation on agar as shown in Figure 1–3.

Several experiments demonstrated the nature of Ab1, Ab2 and Ab3 of such antibodies. It was shown that in the egg can coexist both Ab-1 and Ab-2 as shown in this paper. Both Ab-1 and Ab-2 were separated from the WSF by affinity chromatography. A total of six preparations were made as recorded above. The results of the reproducibility of the indirect ELISA for the determination of anti-anti-idiotypic *Salmonella* antibodies (Ab-3) have high reproducibility as judged by the values of CV.

A new knowledge was gained about the usefulness of the hyperimmune avian egg yolk. It was shown that chicks that had ingested the eggs of chickens immunized with an anti-SpA antibody generated immune responses to *Staphylococcus aureus*. In future work we should purify and characterize anti-*Salmonella* antibodies [5] as well as anti-HIV antibodies in the same manner as we did for anti-SpA antibodies, which were purified and tested against the pathogen. The question is it discovered here widens new perspectives for immunotherapy of existing and emerging infectious diseases? In particular, the preliminary experiments' results open new horizons for future research in the development of oral vaccination for common and neglected microorganisms [17].

Egg yolk constitutes a critical and alternative source of polyclonal antibodies. They present of course some advantages over mammalian antibodies regarding animal welfare, productivity, and specificity [1]. The main antibody (IgY) presents in avian blood (IgY) is transmitted to their offspring and stored at the egg yolks, which facilitates non-invasive harvesting of a large amounts of antibodies [17, 18]. Moreover, due to phylogenetic distance and structural molecular features, IgY is more suitable for treatment and diagnostic purposes than mammalian due to lack of reactivity with human complement system and B cell repertoire [1]. However, IgY displays greater avidity for mammalian conserved polypeptides [15,16]. Indeed, chicken IgY antibodies can be used therapeutically and diagnostically; for example, in the diagnosis of viruses, bacteria, and fungi [19–20]; and the treatment of infectious diseases in both animals and humans [21, 22].

## Conclusions

The present results are encouraging and reflect the chicken’s successful respond of specific antibodies against HIV, *Salmonella* spp. and *S. aureus*.

## Acknowledgements

To the Campus Research and Publication Fund. University of the West Indies, Mona Campus, Jamaica for funding this research.

## Supporting information

**Table A.** Reproducibility (intra- and inter-assay) of the 3 experimental HIV vaccines, including the results for ELISAs for anti-HIV-gp41 (579-601), anti-HIV-gp41 (579 The coefficient of variation intra- and inter-assay were within the normal limits. It suggests that the assays were reproducible. We could not compare our HIV tests with a commercially prepared assay because at the time of this study was not available and it is still not available.

| Vaccines | Avian egg yolk samples tested | Mean Optical Density (OD) | Coefficient of Variation (%) (Intra-assay) | Mean Optical Density (OD) | Coefficient of Variation (%) (Inter-assay) |
| --- | --- | --- | --- | --- | --- |
| HIV-gp41 (579-601) | Positive (n=5)<br>Post-vaccinated | 0.903 | 3.25 | 0.887 | 7.80 |
| HIV-gp41 (579-601) | Negative (n=5)<br>Pre-vaccinated | 0.178 | 3.79 | 0.263 | 6.51 |
| HIV-gp120 (308-331) | Positive (n=5)<br>Post-vaccinated | 0.826 | 4.18 | 0.915 | 8.90 |
| HIV-gp120 (308-331) | Negative (n=5)<br>Pre-vaccinated | 0.204 | 3.89 | 0.190 | 10.05 |
| HIV-gp120 (421-438) | Positive (n=5)<br>Post-vaccinated | 0.920 | 4.82 | 0.934 | 6.98 |
| HIV-gp120 (421-438) | Negative (n=5)<br>Pre-vaccinated | 0.185 | 3.86 | 0.277 | 5.73 |

**Table B.**
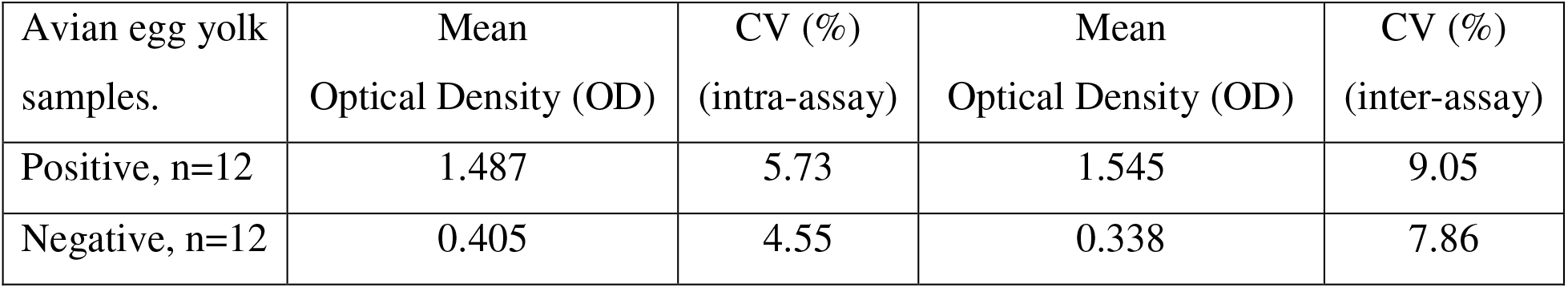
Reproducibility (intra- and inter-assay) of the indirect ELISA for anti-Salmonella antibodies in eggs of brown Leghorn layer hens. The coefficient of variation intra- and inter-assay were within the normal limits. It suggests that the assays were reproducible.

| Avian egg yolk samples. | Mean Optical Density (OD) | CV (%) (intra-assay) | Mean Optical Density (OD) | CV (%) (inter-assay) |
| --- | --- | --- | --- | --- |
| Positive, n=12 | 1.487 | 5.73 | 1.545 | 9.05 |
| Negative, n=12 | 0.405 | 4.55 | 0.338 | 7.86 |

**Table C.** Results of the reproducibility of the indirect ELISA for the determination of anti-anti-idiotypic *Salmonella* antibodies (Ab-3). Positive samples (n=18) were from a group of 18 chicks that were daily fed hyper-immune eggs as stated above. Negative samples were a group from 18 chicks that were daily fed non-hyperimmune eggs and blood was taken at 7, 14 and 28 days to match the previous group. Ab-3 were produced in the sera of the 18 chicks daily fed hyperimmune eggs only.

| Avian purified immunoglobulin Y (IgY) | Mean Optical Density (OD) | CV (%) (intra-assay) | Mean Optical Density (OD) | CV (%) (inter-assay) |
| --- | --- | --- | --- | --- |
| Positive, n=18 | 1.255 | 2.93 | 1.330 | 4.61 |
| Negative, n=18 | 0.319 | 3.07 | 0.378 | 5.78 |

